# Analyzing patterns of tyrosine sulfation in naive antibody repertoires

**DOI:** 10.1101/2022.12.13.520330

**Authors:** Maria Pospelova, Yana Safonova

## Abstract

HIV-1 infects a subset of immune cells identified by the receptor CD4 and a coreceptor, CCR5 or CXCR4. Previous studies revealed bnAbs against HIV-1 with antigen-binding sites mimicking binding sites of CCR5. Such antibodies are characterized by post-translationally sulfated tyrosines and anionic motifs in long complementarity determining regions 3 (CDR3s) of the heavy chains. Despite the great therapeutic potential of human antibodies mimicking CCR5, their immunogenetic signatures remain unknown. In this study, we analyzed human naïve heavy chain antibody repertoires and described the most common VDJ recombination scenarios generating CDR3s with sulfated tyrosines and anionic motifs. We showed ~77% of such CDR3s are generated using seven D genes from two families, IGHD3 and IGHD4. We also demonstrated that sulfated tyrosines and anionic motifs are a common feature of mammalian germline D genes.

## Introduction

### Generation of antibody diversity

Adaptive immune responses are mediated by the production of antibodies (Abs) that bind antigens present on the surfaces of foreign bacteria, viruses, or aberrant cell types. Individual B cells produce antibodies with high specificity to antigens, while broad immunity is provided by a vast array of B cells each producing an antigen-specific antibody. The collection of expressed antibodies in an individual is known as the *antibody repertoire*. The generation of varied antibodies relies on a process called *V(D)J recombination*. This process affects an *immunoglobulin* (*IG*) locus containing the families of the *variable* (*V*), *diversity* (*D*), and *joining* (*J*) genes (*IG genes*) by selecting one V, one D gene, and one J gene, and concatenating them together to generate one of the antibody chains: heavy chain or light chain. Light chain IG loci do not contain D genes and thus encode VJ recombinations. Additionally, VDJ recombination introduces non-genomic insertions, palindromic extensions, and exonuclease removals at the junction of two IG genes. Nucleotides generated using these events are referred to as *junctional diversity*. Each antibody chain includes three antigen-binding loops called *complementarity-determining regions* or *CDRs*. CDR1 and CDR2 are encoded by V gene, and CDR3 covers a short suffix of the V gene, the entire D gene (in case of heavy chain), a prefix of J gene, as well as V-D and D-J junctions (V-J junctions in light chains). Naïve antibodies generated by VDJ recombination are further diversified by gene conversion, class switching, and somatic hypermutations (SHMs) that stochastically improve their binding properties.

### Rational design of HIV vaccines

The most common variant of human immunodeficiency virus HIV-1 infects approximately 1.7 million individuals each year, with an estimated 38 million currently living with the virus. While there are many antiretroviral therapies available, a vaccine would be the most effective measure for controlling the spread of the virus and best hope for eventual eradication. Development of HIV-1 vaccines is an extremely challenging problem due to the high mutation rate of the virus. Strategies of rational HIV vaccine design are aimed at finding antigens that trigger production of broadly neutralizing antibodies (*bnAbs*) that can be generated across the whole population.

### CCR5 mimicry in bnAbs against HIV

HIV-1 infects a subset of immune cells identified by the primary viral receptor CD4 and a coreceptor, usually CCR5 or CXCR4 (Berger et al., 1999). The HIV-1 *envelope* (*Env*) spike protein recognizes the second extracellular loops and the N termini of the coreceptor that are characterized by tyrosines modified by a post-translational addition of sulfate. Antigen binding sites of some bnAbs against HIV-1 acquire sulfated tyrosines thus mimicking CCR5 and binding the Env trimer (Huang et al., 2007). Understanding precise immunogenetic features of precursors of such bnAbs will enable their targeting by vaccines.

### Immunogenetic basis of CCR5-mimicking antibodies

Huttner, 1988 showed that paired amino acids D and E (or *D/E motifs*) are commonly found next to the sulfated tyrosine. The positive training set used by Sulfinator (Monigatti et al., 2002), the state-of-the-art tool for prediction of tyrosine sulfation sites, has abundant D/E motifs around positions of sulfated tyrosines. Huang et al., 2004 demonstrated that, anti-HIV-1 bnAbs with sulfated tyrosine are also characterized by D/E motifs. Another important feature of such bnAbs are long (at least 24 amino acids) CDR3s in heavy chains (Huang et al., 2004; Briney et al., 2012). These observations allow one to identify candidates for bnAbs against HIV-1 in bulk Rep-Seq samples using *in silico* methods and analyze their immunogenetic features. Previous analyses of macaque bnAbs against HIV-1 revealed that CDR3s of their heavy chains are derived from the same common D gene IGHD3-15 (Roark et al., 2021). One amino acid translation of IGHD3-15, YYEDDYGYYYT, contains both D/E motifs and a frequent tyrosine sulfation site at position 6 (underlined). Unlike macaques, none of human germline D genes (and well as fragments of V and J genes contributing to CDR3s) contains D/E motifs. Thus, anionic motifs are either a result of the junctional diversity or SHMs. In this work, we analyzed the role of VDJ recombination events in generation of D/E motifs and sulfated tyrosines in CDR3s of naïve heavy chain antibody repertoires.

## Results

### Finding target CDR3s in naive antibody repertoires

CDR3 sequences from naive heavy chain antibody repertoires collected from 99 donors (PRJEB26509, Gidoni et al., 2019) were extracted and analyzed (see Methods). To find precursors of antibodies that have a potential to mimic CCR5, sulfated tyrosines (further referred to as *sY*) were predicted in CDR3s using Sulfinator, and CDR3s containing both D/E motifs and sY were found. We will refer to sequences containing both D/E motifs and sY as *D/E&sY*. On average, ~6% of CDR3s contain at least one sY (**Figure 1A**) and ~1% of CDR3s contain D/E&sY motifs (**Figure 1B**). Moreover, in ~60% of CDR3s with D/E&sY motifs D/E located next to sY. Such motifs will be referred to as *D/E+sY*. We will call CDR3s containing D/E+sY motifs *target CDR3s*. We noticed that a sY tends to be seen after a D/E motif: D/E+sY motifs are ~9 times more frequent compared to sY+D/E motifs.

### Immunogenetics basis of tyrosine sulfation in human CDR3s

To understand how individual D genes contribute to D/E+sY motifs, CDR3s were aligned to human germline D genes (see Methods). On average, D/E+sY motifs overlap with a D gene in ~86% of target CDR3s and ~7% of CDR3s with sY only. Since no reading frame of any human D gene contains both a D/E motif and sY, a naive target CDR3 has always resulted from the junctional diversity (i.e., specific exonuclease removals and non-genomic insertions). We noticed that D/E+sY overlaps with the start of the D gene 31 times more often than it overlaps with the end of the D gene.

The percentages of aligned CDR3s containing D/E+sY motifs in all aligned CDR3s were calculated for each gene. The mean value is 4%; there are seven genes with the average percentage at least 4%: D4-17, D3-22, D3-10, D3-16, D4-23, D4-11, D3-3 (**Figure 1C**). The most frequent and the second most frequent D genes, D4-17 and IGHD3-22, account for ~34% and ~16% of target CDR3s, respectively. Since all top genes belong to two out of seven human D gene families (IGHD3 and IGHD4), we hypothesize that D genes from the same family use similar scenarios to generate D/E+sY motifs.

Unlike genes D3-22, D3-10, and D3-3, gene D3-16 and all three IGHD4 genes have higher fractions of target CDR3s compared to fractions of CDR3s with sY only. Differences of D3-16 from other IGHD3 genes were discussed below. **Figure 1D** also shows that IGHD3 genes contribute to longer CDR3s compared to IGHD4 genes (P=3.5×10^-205^, the Mann-Whitney U test) thus indicating their possible importance in generation of anti-HIV antibodies.

**Figure 1.**
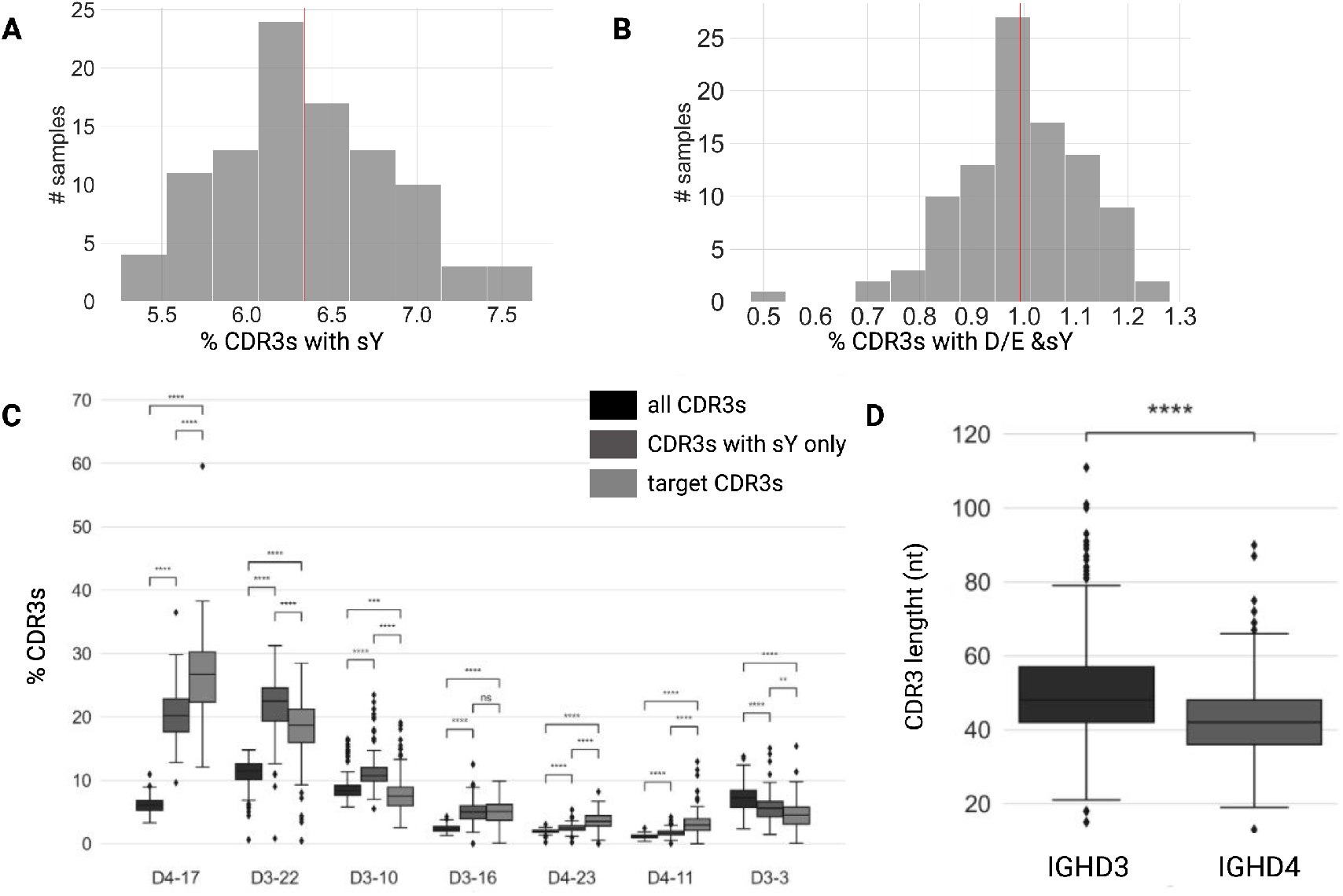
Sulfated tyrosines and D/E motifs in naive antibody repertoires. **(A)** The distribution of fractions of CDR3s with sY with respect to all CDR3s across 99 donors. The red line shows the average value of the distribution. **(B)** The distribution of fractions of CDR3s with D/E&sY motifs with respect to all CDR3s across 99 donors. The red line shows the average value of the distribution. **(C)** D gene usages. Each D gene is shown by three distributions: (i) percentages of CDR3s derived from the D gene (shown as black boxes); (ii) percentages of CDR3s with sY derived from the D gene with respect to all CDR3s with sY (shown as dark gray boxes); and (iii) percentages of target CDR3s derived from the D gene with respect to all target CDR3s. D genes are sorted in the descending order of the average percentage of target CDR3s aligned to them. Only D genes contributing to at least 4% of target CDR3s were shown. Here and further, P-values were computed using the Mann-Whitney U test and denoted as follows: ns:≥0.05; *:<0.05; **:<0.01; ***:<0.001; ****:<0.0001. Boxes show quartiles and the medians of the distributions, outliers are shown as dots and were computed using the interquartile range method implemented in the Python Seaborn package. **(D)** The distribution of nucleotide lengths of CDR3s derived from top seven genes grouped by D gene families IGHD3 and IGHD4 for a single donor (accession number is ERR2567178).

### Unraveling VDJ recombination scenarios in D/E+sY motifs

All seven top D genes generate target CDR3s using their second reading frames. We will refer to VDJ recombination scenarios generating target CDR3s as *target scenarios*. The simplest target scenario involves IGHD4 genes. For example, the second reading frame of the top used D gene D4-17 is DYGDY. **Figure 2A** shows that concatenation of non-genomic nucleotides GA with T, the first nucleotide of D4-17, generated a codon GAT encoding amino acid D. The resulting motif D-D-Y led to sulfation of the tyrosine. **Figure S2** shows that similar target scenarios can be observed for D4-23 and D4-11.

**Figure 2.**
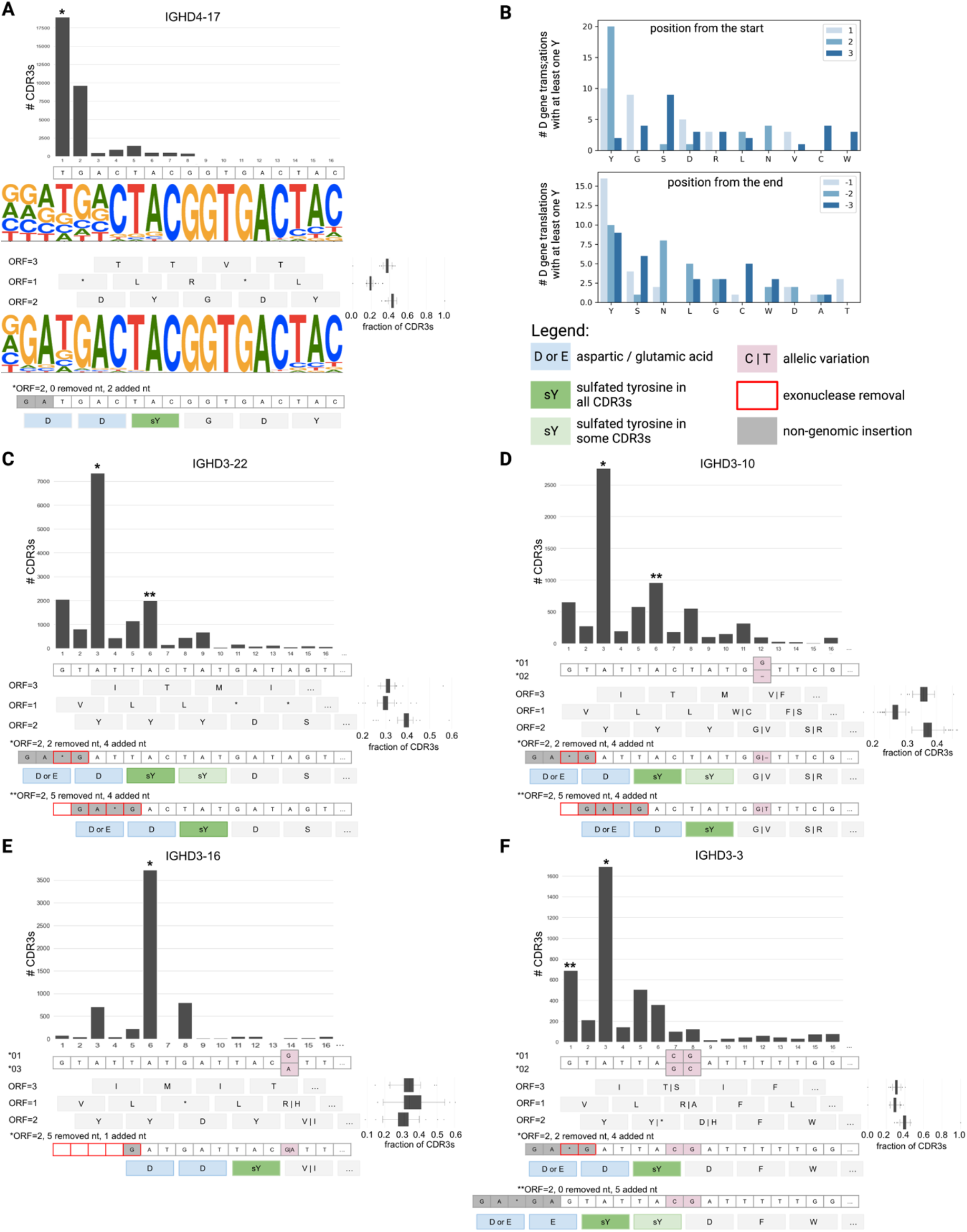
Immunogenetics basis of human heavy chain CDR3s with D/E+sY motifs. **(A)** The target scenario involving gene D4-17 is described by the following plots (from the top to the bottom): (1) the top bar plot shows the total number of target CDR3s where D4-17 gene alignment starts with position *i*, where *i*=1,..,16. The bar(s) corresponding to the target scenario(s) is marked with *(s); (2) the nucleotide sequence of D4-17; (3) the web logo plot shows the alignment of all CDR3 derived from D4-17; (4) three translations of D4-17 and their fractions across 99 donors; (5) the web logo plot shows the alignment of target CDR3s derived from D4-17; (6) the nucleotide sequence corresponding to the target scenario and corresponding amino acid translation. Nucleotides representing the non-genomic insertion are shown as gray boxes. Amino acids D and Y are shown as blue boxes. sYs are shown as bright green boxes. **(B)** The top and bottom bar plots show top 10 amino acids observed at the three first and three last positions of human D gene translations containing at least one Y and the respective counts of D gene translations. Panels **(C), (D), (E),** and **(F)** show target scenarios involving genes D3-22, D3-10, D3-16, and D3-3, respecively. Panel components are consistent with **(A)**, except for web logo plots that were omitted in panels **(C)-(F)**. For all IGHD3 genes, only the first 16 nucleotides are shown. Allele D3-16*03 refers to the novel allele detected by Safonova and Pevzner, 2019. Additional legend: allelic variations at first 16 nt are shown as pink boxes; positions of nucleotides corresponding to exonuclease removals at the 5’ end of the D gene are shown as boxes with red border; tyrosines that are predicted to be sulfated only in a portion of CDR3s corresponding to the target scenario are shown as light green boxes.

**Figure 2B** shows that Y is the most common amino acid observed at the first three and the last three positions of all reading frames of human D genes containing at least one Y. Amino acid D is encoded by codons GAC and GAT that can be obtained from the Y codons (TAC and TAT) by removal of the first nucleotide and addition of a random nucleotide G at the 5’ end of the D gene. Although Y is also the most frequent amino acid located at the 3’ end of D gene translations, it can only be turned into stop codons (TAA and TAG) by removing the last nucleotide and generating a random one. This likely explains the observation that D/E+sY motifs overlap with 5’ ends of D genes more often compared to 3’ ends of D genes.

Analysis of top IGHD3 genes showed that most observed D/E+sY motifs can be explained by the transformation of the first Y into D and completing the motif with D or E generated through non-genomic insertions. Specific target scenarios involving IGHD3 genes are described below. For the sake of simplicity, we will not distinguish non-genomic insertions and palindromic extensions and call them *random insertions* or *random nucleotides*. We will also refer to exonuclease removals of D genes as simply *removals*.

1. The second reading frames of both D3-22 and D3-10 start with YYYD- and sulfation affects the second or the third tyrosines (**Figure 2C,D**).

a. In the case of sulfation of the second tyrosine, the removal of first two germline nucleotides and addition of a random nucleotides GA*G (corresponding to either D or E codon followed by G, * is any nucleotide) resulted in a D/E+sY motif (D-D-sY or E-D-sY) at the 5’ end of the gene.
b. In the case of sulfation of the third tyrosine, four first germline nucleotides were removed. Similarly to the previous scenario, nucleotides GA*G randomly generated before the D gene completed a D/E+sY motif (D-D-sY or E-D-sY).
2. The second reading frame of D3-16 starts with YYDY-. Target CDR3s derived from D3-16 represent a scenario where five first nucleotides of D3-16 were removed at the 5’ end and a randomly generated nucleotide G was concatenated with the truncated gene thus generating the motif D-D-sY (**Figure 2E**).
3. The second reading frame of D3-3 starts with either YYD- (in case of allele *01) or Y*D- (in case of allele *02, * is a stop codon). Sulfated tyrosine predictions were mostly observed for allele *01. The most frequent target scenario generated a D/E+sY motif (D-D-sY or E-D-sY) through removal of two first germline nucleotides and randomly generated insertion GA*G before the D gene (**Figure 2F**). The second most frequent target scenario generated a D/E+sY motif (D-D-sY, E-D-sY, D-D-Y-sY, or E-D-Y-sY) through randomly generated nucleotides GA*GA concatenated with the untruncated sequence of the D3-3.

**Figure S3** shows percentages of target CDR3s corresponding to the described scenarios in naïve antibody repertoires. The most frequent target scenario involving gene D4-17 (GA insertion + 0 removed nt) represents ~0.10% of all CDR3s. Two target scenarios involving gene D3-22 (GA*G insertion + 2 removed nt and GA*G insertion + 5 removed nt) represent ~0.04% and 0.01% of all CDR3s, respectively. The target scenario involving gene D3-16 (G insertion + 5 removed nt) represents ~0.02% of all CDR3s. The remaining target scenarios represent less than 0.01% of all CDR3s each.

The impact of individual alleles of D genes was discussed in **Supplemental Note “The impact of D gene alleles on generation of target CDR3s”**.

### D/E+sY motifs in mammalian germline D genes

To analyze fractions of D/E+sY motifs in mammalian antibodies, Rep-Seq data of various species are needed. Since the availability of non-human antibody repertoires is limited, we analyzed sequences of germline D genes predicted in assemblies of 36 mammalian species representing 11 taxonomic orders from the Vertebrate Genome Project (Rhie et al., 2021). A complete list of species is provided in **Table S1**. IGDetective tool (Sirupurapu et al., 2022) was used to predict sequences of germline D genes, the number of D gene predictions varies from 3 to 104 (**Table S1**).

For each D gene, target scenarios were generated. A target scenario using a D gene is characterized by a random insertion before the D gene and a removal at the 5’ end of the D gene. Using target scenarios illustrated in **Figure 2**, we selected the four common insertions leading to formation of a D/E motif (empty insertion, G, GAGG, GATG) and the removal lengths varying from 0 to 5 nt. The nucleotide sequence corresponding to a target scenario was generated as a concatenation of the insertion and the truncated sequence of the D gene. Each nucleotide sequence resulting from a target scenario was translated into three amino acid sequences, and, for each translation, sY positions were predicted using Sulfinator.

On average, 24% of D genes have at least one target scenario with D/E and sY motifs across 36 species. This suggests that CCR5 mimicry can be used by antibodies of various mammalian species to fight CCR5-binding pathogens (such as lentiviruses). **Figure 3A** shows that there are no clear patterns of D gene fractions associated with the taxonomic orders. **Figure 3B** shows that D genes corresponding to target scenarios have longer lengths compared to D genes without any target scenarios with sY (P=7.2×10^-6^, the Mann-Whitney U test). We classified a 7 aa fragment containing a sY as *germline* if it is fully encoded by D gene and *partial* if it overlaps with a random insertion. **Figure 3C** shows that, as expected, germline motifs are more diverse compared to partial motifs. While partial motifs were detected for all species, germline motifs were found only in 12 species (Asiatic elephant, black rhinoceros, California sea lion, Canada lynx, Chinese rufous horseshoe bat, gray squirrel, greater horseshoe bat, North American porcupine, Pallas’s mastiff bat, platypus, short-beaked echidna, yellow-spotted hyrax) representing 7 orders (**Figure 3D**).

**Figure 3.**
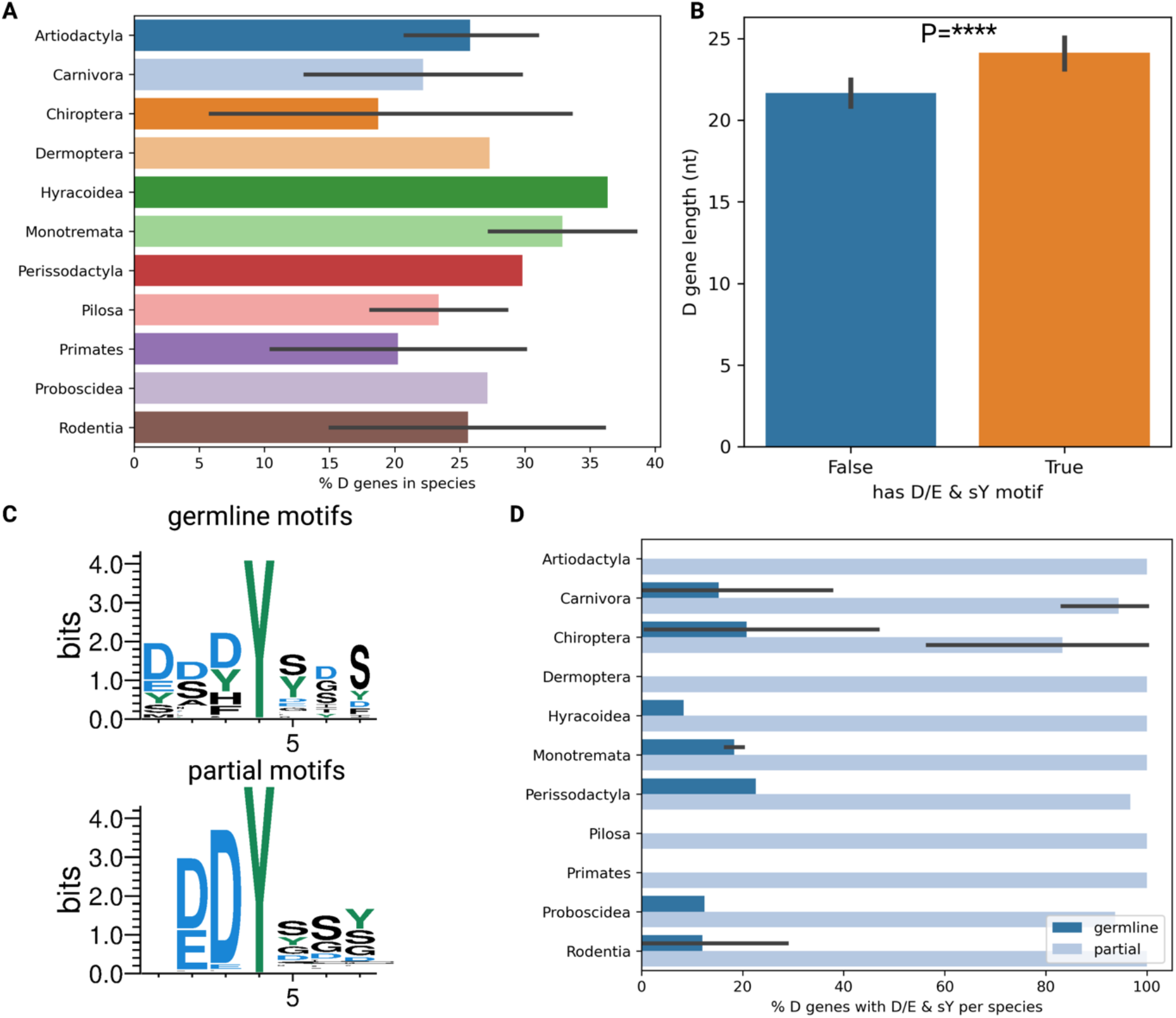
D/E and sY motifs in mammalian D genes. **(A)** The distribution of percentages of D genes with at least one target scenario across 36 species grouped by 11 orders. Here and below, error bars show 95% confidence intervals. **(B)** The distributions of nucleotide lengths of D genes grouped by whether they have at least one target scenario. **(C)** Web logo plots showing partial and germline motifs. sY are shown at position 4 in each plot. **(D)** The distribution of percentages of D genes containing partial and germline D/E&sY motifs across 36 species grouped by 11 orders.

## Discussion

In this study, patterns of tyrosine sulfation were analyzed in complementarity-determining regions 3 of human naïve heavy chain antibody repertoires collected across 99 donors. For each donor, CDR3s containing both predicted sites of sulfated tyrosines (sY) and anionic motifs (D/E motifs) were identified. We showed that, in the most common configurations, a D/E motif is located right before a sY. We referred to such motifs as D/E+sY. We also showed that D/E+sY motifs overlap with D genes in 86% of cases. While the sY is always fully encoded by the D gene, the D/E motif is a joint result of exonuclease removals at the 5’ end of the D gene and non-genomic insertions. Despite the fact that none of human D genes fully encodes a D/E+sY motif, we revealed strong associations between the motif and two D gene families: IGHD3 and IGHD4. We demonstrated that seven genes from these two families (IGHD4-17, IGHD3-22, IGHD3-10, IGHD3-16, IGHD4-23, IGHD4-11, and IGHD3-3) account for ~77% of CDR3s with D/E+sY motifs. IGHD3 genes contribute to longer CDR3s compared to IGHD4 genes thus indicating their possible importance for generation of HIV-specific antibodies.

The second reading frames of four revealed IGHD3 genes start with YY. The most common VDJ recombination scenarios generating D/E+sY motifs include the transformation of the first Y codon into a D codon (through the truncation of the first nucleotide of the D gene and generation of a random nucleotide G) that, in combination with a D or E codon generated through non-genomic insertions, leads to sulfation of the second Y (encoded by the D gene). The second reading frames of three IGHD4 genes start with DY. VDJ recombination scenarios involving IGHD4 genes concatenate a randomly generated D or E codon with the D gene leading to sulfation of the tyrosine encoded by the D gene. We showed that the most common VDJ recombination scenarios generating D/E+sY motifs use IGHD4-23, IGHD3-22, and IGHD3-16 genes.

The analysis of germline D genes predicted in genomes of 36 mammalian species showed that similar VDJ recombination scenarios are possible for ~24% of D gene per species. Moreover, D/E motifs and predicted tyrosine sulfation sites are encoded in the germline D genes of 12 out of 36 species. Since D genes are extremely diverse (very few D genes are shared among two or more species), we assume that D genes (fully or partially) encoding D/E+sY motifs might be a subject of the selective pressure. We also hypothesize that these D genes can generate antibodies fighting CCR5-binding pathogens (such as lentiviruses). Further analysis of non-human antibody repertoires (both naïve and antigen-responsive) will shed light on the role of these D genes

## Methods

### Repertoire sequencing data preprocessing

Rep-Seq reads were aligned to V and J genes using the Diversity Analyzer tool (Shlemov et al., 2017); sequences aligned to light chain IG genes were discarded. CDR3s reported by the Diversity Analyzer tool were translated into amino acid sequences. CDR3s with lengths below 5 amino acids were discarded. Positions of sulfated tyrosines were predicted using Sulfinator with the default E-value threshold (Monigatti et al., 2002). Sample ERR2567275 contains only 6 distinct CDR3s and was discarded from further analysis.

### Aligning germline D genes to naïve CDR3s

For each CDR3, D genes with at least 7 nt long matches were selected. A CDR3 was aligned to a D gene with a match *M* if the following conditions were met: (i) *M* is the longest match across all other matches; (ii) there are no other D genes with matches of the same length. If two or more longest matches from two or more D genes were found, the corresponding CDR3 was considered unaligned and was excluded from the further analysis. D gene alleles were treated as sequences of the same gene.

## Notes

### Competing Interest Statement

The authors have declared no competing interest.

### Summary of Updates

minor fixes

