## Supplemental Materials for "Analyzing patterns of tyrosine sulfation in naive antibody repertoires"

Supplemental Materials
for

### Analyzing patterns of tyrosine sulfation in naive antibody repertoires

Maria Pospelova [1] and Yana Safonova [2,*]
1 Faculty of IT and Programming, ITMO University, Saint Petersburg, Russia
2 Department of Computer Science, Johns Hopkins University, Baltimore, USA

include 3 Supplemental Figures, 1 Supplemental Table, and 1 Supplemental Note.

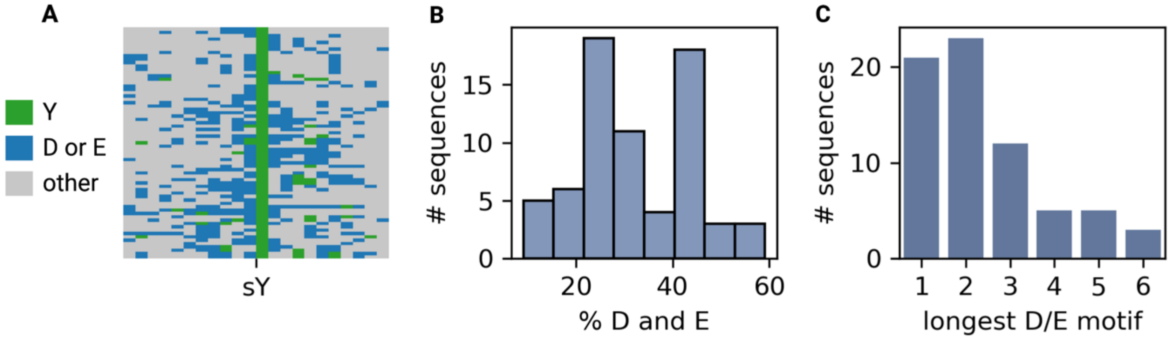


**Figure S1. The positive training set of Sulfinator consists of sequences with D/E motifs. (A)** Alignment is 22 aa long fragments of protein sequences used as the positive training set. The sulfated tyrosine is shown in the middle as the green column. **(B)** The percentages of amino acids D and E in sequences from the positive training set. **(C)** Lengths of the longest motif containing amino acids D and E only in each sequence from the positive training set.

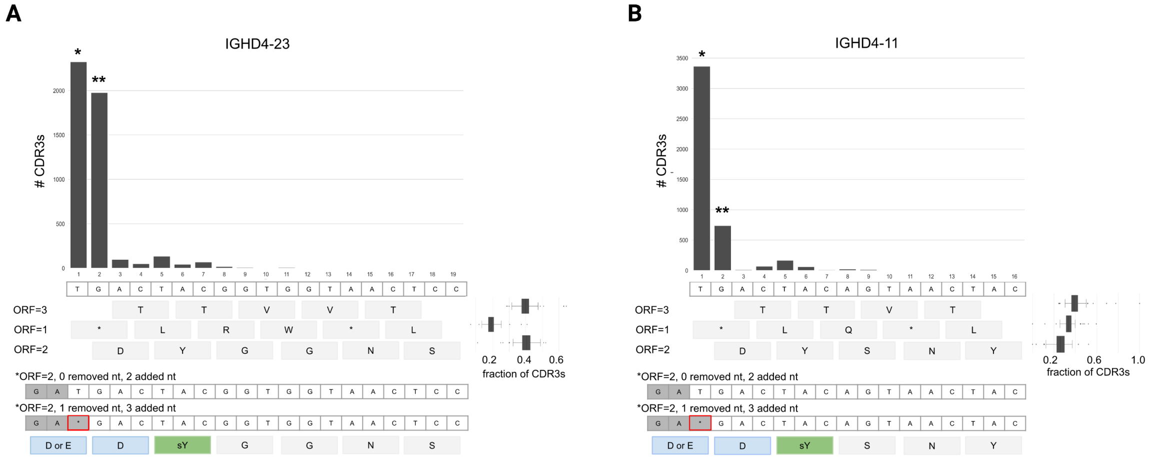


**Figure S2. Target scenarios using D genes D4-23 (A) and D4-11 (B).** The legend is described in the caption to **Figure 2.**

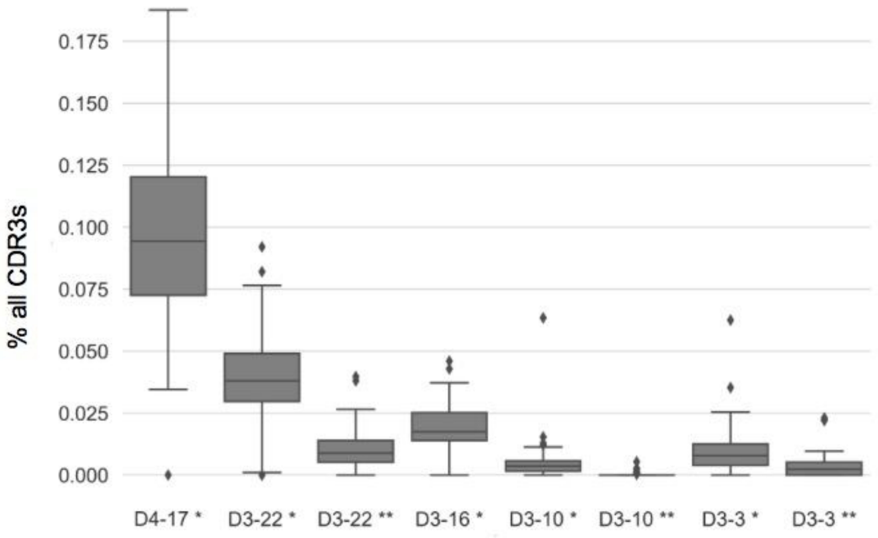


**Figure S3. Percentages of CDR3s representing target scenarios in Figure 2 across 99 donors.** Target scenarios are labeled with gene name and *s, labels are consistent with **Figure 2.**

| # | Order | Latin name | Common name | Accession number | # D gene predictions |
|---|---|---|---|---|---|
| 1 | Artiodactyla | Balaenoptera musculus | blue whale | mBalMus1 | 4 |
| 2 | Artiodactyla | Cervus elaphus | red deer | mCerEla1 | 11 |
| 3 | Artiodactyla | Mesoplodon densirostris | Blainville's beaked whale | mMesDen1 | 5 |
| 4 | Artiodactyla | Phocoena sinus | vaquita | mPhoSin1 | 3 |
| 5 | Artiodactyla | Orcinus orca | killer whale | mOrcOrc1 | 5 |
| 6 | Artiodactyla | Tursiops truncatus | common bottlenose dolphin | mTurTru1 | 5 |
| 7 | Carnivora | Canis lupus orion | Greenland wolf | mCanLor1 | 7 |
| 8 | Carnivora | Lutra lutra | Eurasian river otter | mLutLut1 | 17 |
| 9 | Carnivora | Lynx canadensis | Canada lynx | mLynCan4 | 9 |
| 10 | Carnivora | Mustela erminea | stoat | mMusErm1 | 9 |
| 11 | Carnivora | Neofelis nebulosa | clouded leopard | mNeoNeb1 | 4 |
| 12 | Carnivora | Zalophus californianus | California sea lion | mZalCal1 | 15 |
| 13 | Chiroptera | Molossus molossus | Pallas's mastiff bat | mMolMol1 | 46 |
| 14 | Chiroptera | Myotis myotis | greater mouse-eared bat | mMyoMyo1 | 3 |
| 15 | Chiroptera | Phyllostomus discolor | pale spear-nosed bat | mPhyDis1 | 12 |
| 16 | Chiroptera | Pipistrellus kuhlii | Kuhl's pipistrelle | mPipKuh1 | 16 |
| 17 | Chiroptera | Pipistrellus pipistrellus | common pipistrelle | mPipPip1 | 21 |
| 18 | Chiroptera | Rhinolophus ferrumequinum | greater horseshoe bat | mRhiFer1 | 11 |
| 19 | Chiroptera | Rhinolophus sinicus | Chinese rufous horseshoe bat | mRhiSin1 | 19 |
| 20 | Dermoptera | Cynocephalus volans | Philippine flying lemur | mCynVol1 | 22 |
| 21 | Hyracoidea | Heterohyrax brucei | yellow-spotted hyrax | mHetBru1 | 33 |
| 22 | Monotremata | Ornithorhynchus anatinus | platypus | mOrnAna1 | 22 |
| 23 | Monotremata | Tachyglossus aculeatus | short-beaked echidna | mTacAcu1 | 13 |
| 24 | Perissodactyla | Diceros bicornis | black rhinoceros | mDicBic1 | 104 |
| 25 | Pilosa | Choloepus didactylus | two-toed sloth | mChoDid1 | 21 |
| 26 | Pilosa | Tamandua tetradactyla | southern tamandua | mTamTet1 | 22 |
| 27 | Primates | Callithrix jacchus | common marmoset | mCalJac1 | 19 |
| 28 | Primates | Lemur catta | ring-tailed lemur | mLemCat1 | 10 |
| 29 | Proboscidea | Elephas maximus | Asiatic elephant | mEleMax1 | 59 |
| 30 | Rodentia | Arvicola amphibius | European water vole | mArvAmp1 | 8 |
| 31 | Rodentia | Arvicanthis niloticus | African grass rat | mArvNil1 | 38 |
| 32 | Rodentia | Erethizon dorsatum | North American porcupine | mEreDor1 | 27 |
| 33 | Rodentia | Jaculus jaculus | lesser Egyptian jerboa | mJacJac1 | 7 |
| 34 | Rodentia | Rattus norvegicus | Norway rat | mRatNor1 | 33 |
| 35 | Rodentia | Sciurus carolinensis | gray squirrel | mSciCar1 | 9 |
| 36 | Rodentia | Sciurus vulgaris | Eurasian red squirrel | mSciVul1 | 4 |

**Table S1. Mammalian species used for detection of germline D genes.** Accession numbers are consistent with IDs used by the Vertebrate Genome Project. Gray and white lines separate species from different orders.

#### Supplemental Note “The impact of D gene alleles on generation of target CDR3s”

Three top D genes (D3-10, D3-16, and D3-3) have more than one known allele. To analyze their impact on generation of target CDR3s, each CDR3 derived from one of these three D genes was assigned to one of known alleles. If two or more alleles were aligned with the same score, the corresponding CDR3 was classified as *undefined*. For gene D3-16, the novel allele with nucleotide sequence GTATTATGATTACATTTGGGGGAGTTATCGTTATACC detected by Safonova and Pevzner, 2019 was included. We referred to this allele as IGHD3-16*03.

On average, ~94.7% and ~0.7% of CDR3s derived from D3-10 were aligned to alleles *01 and *02, respectively (**Figure S4A**). We thus assumed that most individuals were homozygous by allele *01. As the result, the majority of target CDR3s were derived from D3-10*01 as well. To make conclusions about usage of allele D3-10*02 in target CDR3s a larger and more diverse cohort is needed.

In case of D3-3, ~63.5% and ~0.2% of CDR3s were aligned to alleles *01 and *02, respectively, and for ~36.3% of CDR3s, the allele could not be determined (**Figure S4B**). The average percentage of target CDR3s derived from D3-3 and aligned to allele *01 (*02) is ~75.2% (<1%). We assume that low usages of allele *02 can be explained by the presence of stop codon in the second ORF.

On average, ~52.1% of CDR3s derived from D3-16 were aligned to allele *02 (**Figure S4C**). Percentages of CDR3s aligned to alleles *01 and *03 are ~4.0 and ~12.4, respectively. Alignments of target CDR3s followed the same trends: ~6.7%, ~32.1%, and ~9.4% of target CDR3s derived from D3-16 were aligned to alleles *01, *02, and *03, respectively.

In summary, the analyzed cohort could not provide us with sufficient information to make conclusions about the role of alleles of D3-10, D3-3, and D3-16. For all three D genes, the most popular allele contributed more to production of target CDR3s compared to other alleles. For genes D3-3 and D3-16, the analysis was complicated by high percentages of CDR3s that could not be assign to an allele.

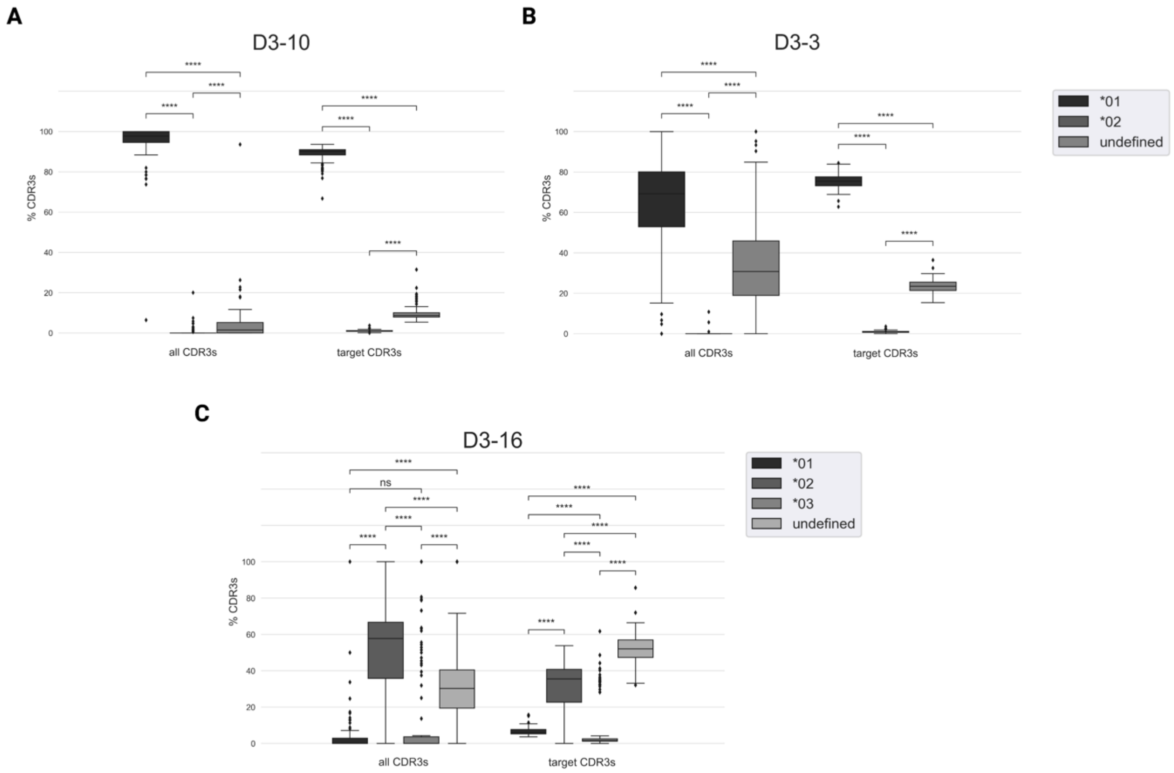


**Figure S4. The percentages of all CDR3s and target CDR3s derived from D genes D3-10 (A), D3-3 (B), and D3-16 (C) and aligned to their alleles.**
